# Auto-CORPus: A Natural Language Processing Tool for Standardising and Reusing Biomedical Literature

**DOI:** 10.1101/2021.01.08.425887

**Authors:** Tim Beck, Tom Shorter, Yan Hu, Zhuoyu Li, Shujian Sun, Casiana M. Popovici, Nicholas A. R. McQuibban, Filip Makraduli, Cheng S. Yeung, Thomas Rowlands, Joram M. Posma

## Abstract

To analyse large corpora using machine learning and other Natural Language Processing (NLP) algorithms, the corpora need to be standardised. The BioC format is a community-driven simple data structure for sharing text and annotations, however there is limited access to biomedical literature in BioC format and a lack of bioinformatics tools to convert online publication HTML formats to BioC. We present Auto-CORPus (Automated pipeline for Consistent Outputs from Research Publications), a novel NLP tool for the standardisation and conversion of publication HTML and table image files to three convenient machine-interpretable outputs to support biomedical text analytics. Firstly, Auto-CORPus can be configured to convert HTML from various publication sources to BioC. To standardise the description of heterogenous publication sections, the Information Artifact Ontology is used to annotate each section within the BioC output. Secondly, Auto-CORPus transforms publication tables to a JSON format to store, exchange and annotate table data between text analytics systems. The BioC specification does not include a data structure for representing publication table data, so we present a JSON format for sharing table content and metadata. Inline tables within full-text HTML files and linked tables within separate HTML files are processed and converted to machine-interpretable table JSON format. Finally, Auto-CORPus extracts abbreviations declared within publication text and provides an abbreviations JSON output that relates an abbreviation with the full definition. This abbreviation collection supports text mining tasks such as named entity recognition by including abbreviations unique to individual publications that are not contained within standard bio-ontologies and dictionaries.

**Availability:** The Auto-CORPus package is freely available with detailed instructions from Github at https://github.com/omicsNLP/Auto-CORPus/.

## Introduction

Natural language processing (NLP) is a branch of artificial intelligence that uses computers to process, understand and use human language. NLP is applied in many different fields including language modelling, speech recognition, text mining and translation systems. In the biomedical realm NLP has been applied to extract, for example, medication data from electronic health records and patient clinical history from free-text (unstructured) clinical notes, to significantly speed up processes that would otherwise be extracted manually by experts (1, 2). Biomedical research publications, although semi-structured, pose similar challenges with regards to extracting and integrating relevant information (3). The full-text of biomedical literature is predominately made available online in the accessible and reusable Hypertext Markup Language (HTML) format, however, some publications are only available as Portable Document Format (PDF) documents which are more difficult to reuse. Efforts to resolve the problem of publication text accessibility across science in general includes work by the Semantic Scholar search engine to convert PDF documents to HTML formats (4). Whichever process is used to obtain a suitable HTML file, before the text can be processed using NLP, heterogeneously structured HTML requires standardisation and optimisation. BioC is a simple JSON (and XML) format for sharing and reusing text data that has been developed by the text mining community to improve system interoperability (5). The BioC data model consists of collections of documents divided into data elements such as publication sections and associated entity and relation annotations. PubMed Central (PMC) makes full-text articles from its Open Access and Author Manuscript collections available in BioC format (6). To our knowledge there are no services available to convert PMC publications that are not part of these collections to BioC. Additionally, there is a gap in available software to convert publishers’ publication HTML to BioC, creating a bottleneck in many biomedical literature text mining workflows caused by having to process documents in heterogenous formats. To bridge this gap, we have developed an Automated pipeline for Consistent Outputs from Research Publications (Auto-CORPus) that can be configured to process any HTML publication structure and transform the corresponding publications to BioC format.

During information extraction, the publication section context of an entity will assist with entity prioritisation. For example, an entity identified in the *results section* may be regarded as a higher priority novel finding than one identified in the *introduction section*. However, the naming and the sequential order of sections within research articles differ between publications. A *methods section*, for example, may be found at different locations relative to other sections and identified using a range of synonyms such as *experimental section*, *experimental procedures* and *methodology*. The Information Artifact Ontology (IAO) was created to serve as a domain-neutral resource for the representation of types of information content entities such as documents, databases, and digital images (7). Auto-CORPus applies IAO annotations to BioC file outputs to standardise the description of sections across all processed publications.

Vast amounts of biomedical data are contained in publication tables which can be large and multi-dimensional where information beyond a standard two-dimensional matrix is conveyed to a human reader. For example, a table may have subsections or entirely new column headers to merge multiple tables into a single structure. Milosevic and colleagues developed a methodology to analyse complex tables that are represented in XML format and perform a semantic analysis to classify the data types used within a table (8). The outputs from the table analysis are stored in esoteric XML or database models. The communal BioC format on the other hand has limited support for tables, for example the PMC BioC JSON output includes table data in PMC XML format, introducing file parsing complexity. In addition to variations in how tables are structured, there is variability amongst table file types. Whereas publication full-text is contained within a single HTML file, tables may be contained within that full-text file (inline tables), or individual tables may be contained in separate HTML files (linked tables). We have defined a dedicated table JSON format for representing table data from both formats of table. The contents of individual cells are unambiguously identified and thus can be used in entity and relation annotations. In developing the Auto-CORPus table JSON format, we adopted a similar goal to the BioC community, namely, a simple format to maximise interoperability and reuse of table documents and annotations. The table JSON reuses the BioC data model for entity and relation annotations, ensuing that table and full-text annotations can share the same BioC syntax. Auto-CORPus transforms both inline and linked HTML tables to the machine interpretable table JSON format.

Abbreviations and acronyms are widely used in publication text to reduce space and avoid prolix. Abbreviations and their definitions are useful in text mining to identify lexical variations of words describing identical entities. However, the frequent use of novel abbreviations in texts presents a challenge for the curators of biomedical lexical ontologies to ensure they are continually updated. Several algorithms have been developed to extract abbreviations and their definitions from biomedical text (9–11). Abbreviations within publications can be defined when they are declared within the full-text, and in some publications, are included in a dedicated *abbreviations section*. Auto-CORPus adapts an abbreviation detecting methodology (12) and couples it with IAO section detection to comprehensively extract abbreviations declared in the full-text and in the *abbreviations section*. For each publication, Auto-CORPus generates an abbreviations dictionary JSON file.

The aim of this paper is to describe the open Auto-CORPus python package and the text mining use cases that make it a simple user-friendly application to create machine interpretable biomedical literature files, from a single publication to a large corpus. The following sections describe the technical details about the algorithms developed and the bench-marking undertaken to assess the quality of the three Auto-CORPus files generated for each publication: BioC full-text, Auto-CORPus tables and abbreviations JSONs.

## Materials and Methods

### Data for algorithm development

We used a set of 3,279 full-text HTML and 1,041 linked table files to develop and test the algorithms described in this section. Files for 1,200 Open Access (OA) Genome-Wide Association Study (GWAS) publications whose data exists in the GWAS Central database (13) were downloaded from PMC in March 2020. A further 1,241 OA PMC publications of Metabolome-Wide Association Studies (MWAS) and metabolomics studies on cancer, gastrointestinal diseases, metabolic syndrome, sepsis and neurodegenerative, psychiatric, and brain illnesses were also downloaded to ensure the methods are not biased towards one domain, more information is available in the Supplementary Materials. This formed a collection of 2,441 publications that will be referred to as the “OA dataset”. We also downloaded publisher-specific full-text files, and linked table data where available, for publications whose data exists in the GWAS Central database. This collection of 838 full-text and 1,041 table HTML files will be referred to as the “publisher dataset”. Table 1 lists the publishers and journals included in the publisher dataset and the number of publications that overlap with the OA dataset.

**Table 1.** Publishers and journals included in the publisher dataset. The full-text files were downloaded in HTML format and the linked table files were downloaded when available in HTML formats. The full-text files that overlap with the OA dataset were used to assess the consistency of outputs generated from different sources.

| **Publisher** | **Journal(s)** | **Number of full-text files** | **Overlap with OA dataset** | **Table type** | **Number of table files** |
| --- | --- | --- | --- | --- | --- |
| American Heart Association | Circulation<br>Cardiovascular Genetics | 52 | 39 | Inline | - |
| American Society of Hematology | Blood | 31 | 25 | Inline | - |
| American Thoracic Society | American Journal of Respiratory and Critical Care Medicine | 20 | 18 | Inline | - |
| BioMed Central | BMC Medical Genetics | 43 | 43 | Linked HTML | 160 |
| Cell Press | American Journal of Human Genetics | 5 | 5 | Inline | - |
| Elsevier | Biological Psychiatry | 5 | 5 | Inline | - |
| Elsevier | Gastroenterology | 5 | 2 | Inline | - |
| Frontiers | Frontiers in Genetics | 20 | 20 | Linked images | n/a |
| Massachusetts Medical Society | The New England Journal of Medicine | 20 | 12 | Linked images | n/a |
| Mosby | The Journal of Allergy and Clinical Immunology | 5 | 3 | Inline | - |
| Nature Portfolio | European Journal of Human Genetics | 50 | 50 | Linked HTML | 123 |
| Nature Portfolio | Journal of Human Genetics | 37 | 3 | Linked HTML | 90 |
| Nature Portfolio | Molecular Psychiatry | 103 | 78 | Linked HTML | 262 |
| Nature Portfolio | Scientific Reports | 80 | 80 | Linked HTML | 190 |
| Nature Portfolio | The Pharmacogenomics Journal | 37 | 16 | Linked HTML | 116 |
| Nature Portfolio | Translational Psychiatry | 41 | 41 | Linked HTML | 87 |
| Oxford University Press | Human Molecular Genetics | 254 | 186 | Inline | - |
| PLOS | PLOS One | 20 | 20 | Linked images | n/a |
| Springer | Human Genetics | 5 | 2 | Linked HTML | 13 |
| Wiley-Blackwell | American Journal of Medical Genetics | 5 | 0 | Inline | - |
| **Total** |  | **838** | **648** |  | **1041** |

#### PubMed Central (PMC) search terms

There exists no equivalent to the GWAS Central database (13) for MWAS/metabolomics studies, therefore a best effort was made to create a collection of publications that span multiple publishers, journals and focus on a variety of disease traits. We searched PMC in March 2020 for OA publications using the following search terms to appear in the title or abstract for the field (metabolomics, metabolome, metabolomic, metabolome-wide, metabonomics, metabonome, metabonomic, metabonome-wide, lipidomics, lipidome, lipidomic, lipidome-wide, metabolites, metabolite, metabolic profiling, metabolic phenotyping), type of samples (urine, urinary, blood, serum, plasma, faecal, faeces, cerebrospinal fluid, CSF, biofluid, stool, feces, fecal), type of technology (nmr spectroscopy, nuclear magnetic resonance, NMR, mass spectrometry, LCMS, GCMS, UPLCMS, LC-MS, GCMS, UPLC-MS, CEMS, CE-MS), human studies (human, patients, subjects, participants) – all using an ‘AND’ statement – and exclude terms (using a ‘NOT’ statement) outside the scope (mouse, mice, rat, rats, dog, dogs, animal, cell culture, dose, review, proteomics, diet, proteomic, proteome, transcriptomics, transcriptomic, transcriptome).

These search terms were combined with either search terms targeting specific journals that commonly publish MWAS/metabolomics studies (Analytical Chemistry, Journal of Proteome Research, Analytica Chimica Acta, Journal of Chromatography A, Metabolites, Scientific Reports, PLOS One, Analytical and Bioanalytical Chemistry, Metabolomics, Proceedings of the National Academy of Sciences of the United States of America) or with specific disease trait terms in the title/abstract. For example, for cancer the additional search terms consist of: cancer, tumor, tumour, cancerous, carcinogen, carcinogenic, carcinoma, leukaemia, leukemia, leukaemic, leukemic, lymphoma, malignancy, premalignancy, pre-malignancy, melanoma, metastasis, sarcoma, adjuvant, neoadjuvant, chemotherapy, chemo therapy, chemotherapy, malignant, premalignant, pre-malignant, precancerous, pre-cancerous, adenocarcinoma, metastatic. The same approach was used for publications relating to gastrointestinal diseases, metabolic syndrome, sepsis and, neurodegenerative, psychiatric, and brain illnesses.

The full-text OA publications were downloaded in HTML format after duplicates (e.g. a ‘cancer’ study from one of the targeted journals) were removed. This resulted in a total of 1,241 unique publications that are included in the OA dataset.

### Algorithms for processing publication full-text HTML

An Auto-CORPus configuration file is set by the user to define the heading and paragraph HTML elements used in the publication files to be processed. Regular expressions can be used within the configuration file allowing a group of publications with a similar but not an identical structure to be defined by a single configuration fil e, for example when processing publications from journals by the same publisher. The heading elements are used to delineate the content of the publication sections and the BioC data structure is populated with publication text. All HTML tags including text formatting (e.g., emphasised words, superscript and subscript) are removed from the publication text. Each section is automatically annotated using IAO (see below) and the BioC data structure is output in JSON format. The BioC specification requires “ key files” to accompany BioC data files to specify how the data files should be interpreted (5). We provide key files to define the da ta elements in the Auto-CORPus JSON output files for full-text, tables, and abbreviations (https://github.com/omicsNLP/Auto-CORPus/tree/main/keyFiles). Figure 1 gives an example of the BioC JSON output and the abbreviations and tables outputs are described below.

**Fig. 1.**
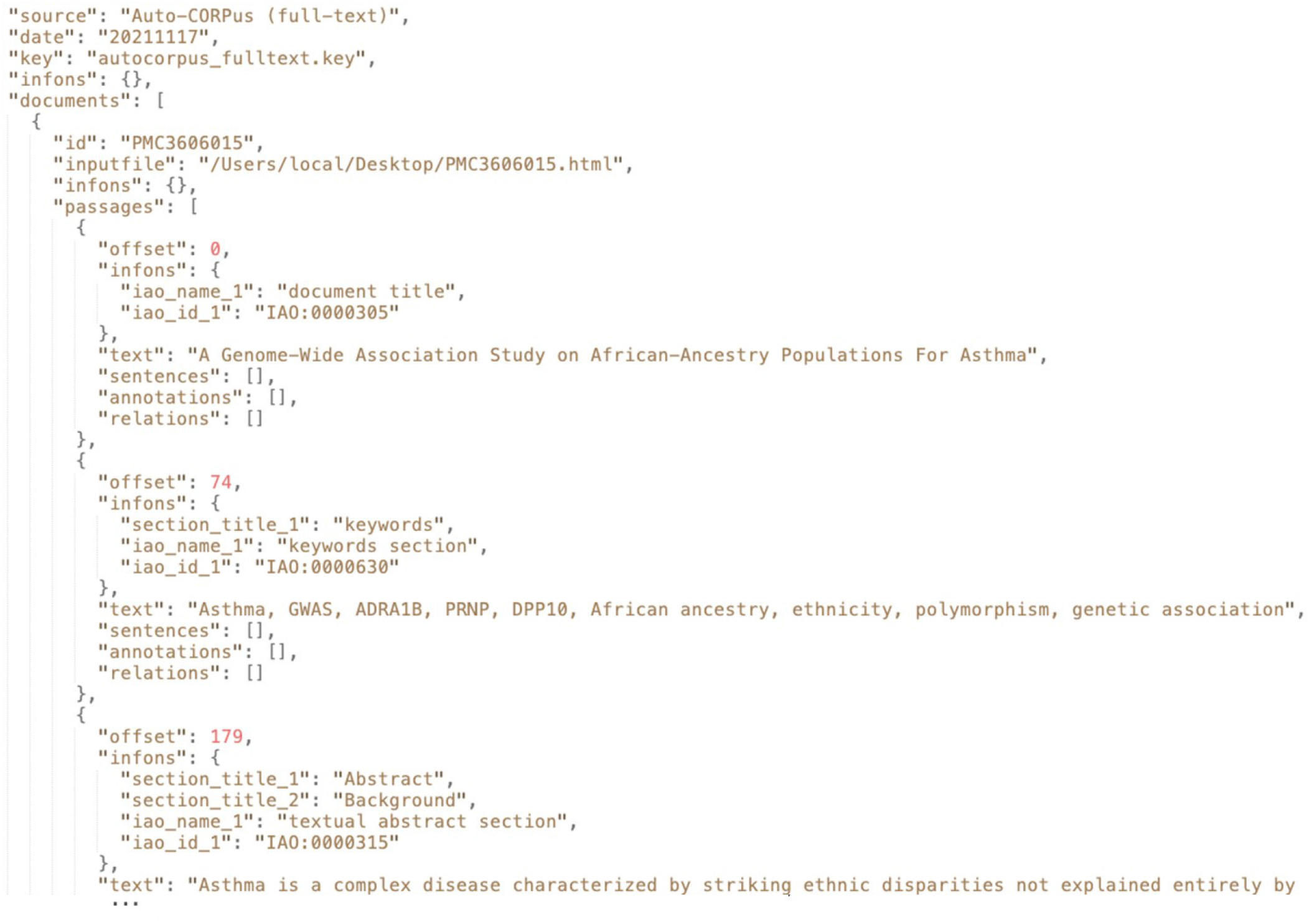
An extract of the Auto-CORPus BioC JSON created from the PMC3606015 (14) full-text HTML file. Each section is annotated with IAO terms. The “autocor-pus_fulltext.key” file describes the contents of the full-text JSON file (https://github.com/omicsNLP/Auto-CORPus/blob/main/keyFiles/autocorpus_fulltext.key).

Abbreviations in the full-text are found using an adaptation of a previously published methodology and implementation (12). The method finds all brackets within a publication and if there are two or more non-digit characters within brackets it considers if the string in the brackets could be an abbreviation. It searches for the characters present in the brackets in the text on either side of the brackets one by one. The first character of one of these words must contain the first character within the bracket, and the other characters within that bracket must be contained by other words that follow the first word whose first character is the same as the first character in that bracket. An example of the Auto-CORPus abbreviations JSON is given in Figure 2 which shows that the output from this algorithm is stored along with the abbreviations defined in the publication abbreviations section (if present).

**Fig. 2.**
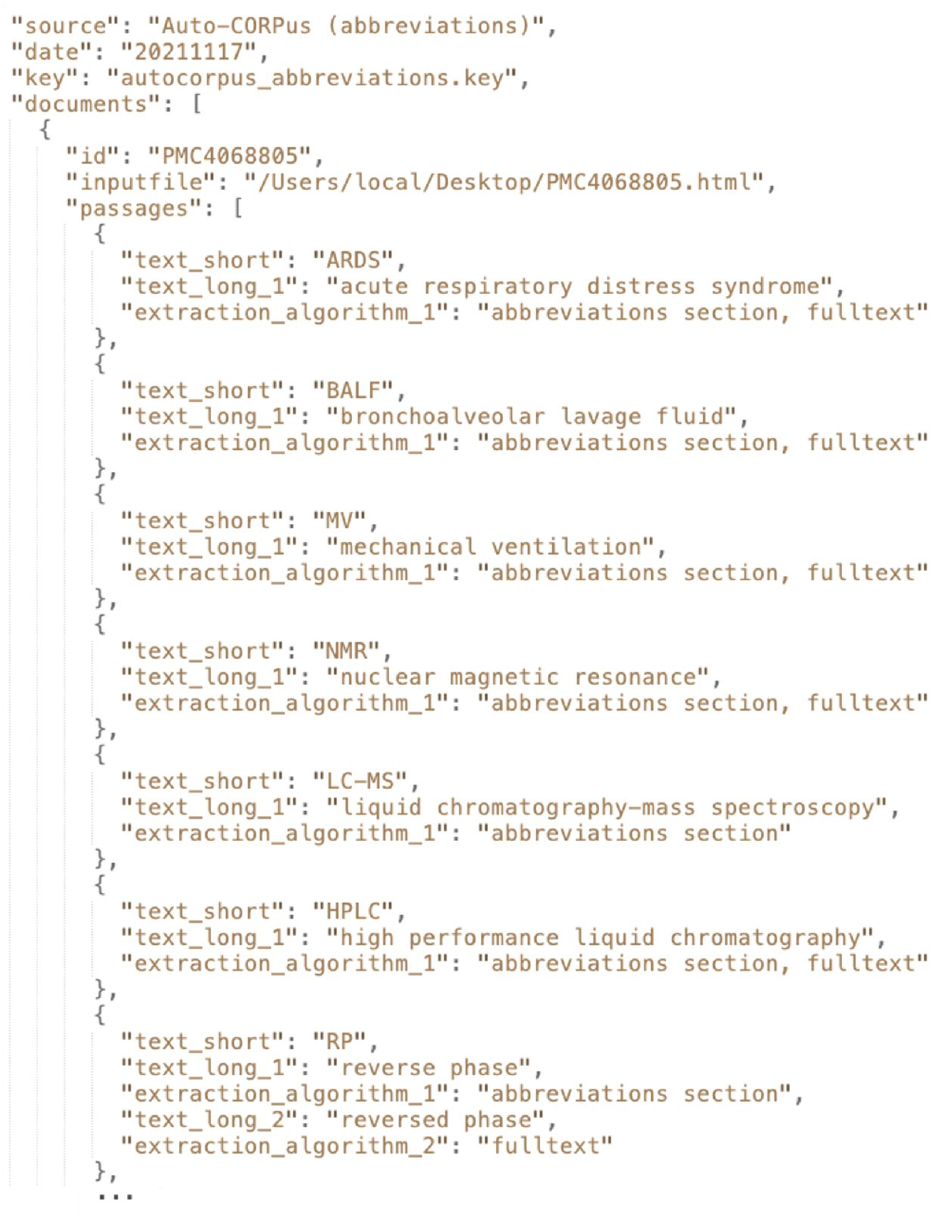
An extract from the Auto-CORPus abbreviations JSON created from the PMC4068805 (15) full-text HTML file. For each abbreviation the corresponding long form definition is given along with the algorithm(s) used to detect the abbreviation. Most of the abbreviations shown were independently identified in both the full-text and in the abbreviations section of the publication. A variation in the definition of “RP” was detected: in the abbreviations section this was defined as “reverse phase”, however in the full-text this was defined as “reversed phase”. The “autocorpus_abbreviations.key” file describes the contents of the abbreviations JSON file (https://github.com/omicsNLP/Auto-CORPus/blob/main/keyFiles/autocorpus_abbreviations.key).

### Algorithms for classifying publication sections with IAO terms

A total of 21,849 section headers were extracted from the OA dataset and directed acyclic graphs (DAGs) were created for each publication (Figure 3). The individual DAGs were then combined into a directed graph (digraph) and the extracted section headers were mapped to IAO (v2020-06-10) document part terms using the Lexical OWL Ontology Matcher (LOOM) method (16). Fuzzy matching using the fuzzywuzzy python package (v0.17.0) was then used to map headers to the preferred section header terms and synonyms, with a similarity threshold of 0.8 (e.g., the typographical error ‘experemintal section’ in PMC4286171 (15) is correctly mapped to *methods section*). This threshold was evaluated by two independent researchers who confirmed all matches for the OA dataset were accurate. Digraphs consist of nodes (entities, headers) and edges (links between nodes) and the weight of the nodes and edges is proportional to the number of publications in which these are found. Here the digraph consists of 372 unique nodes and 806 directed edges (Figure 4).

**Fig. 3.**
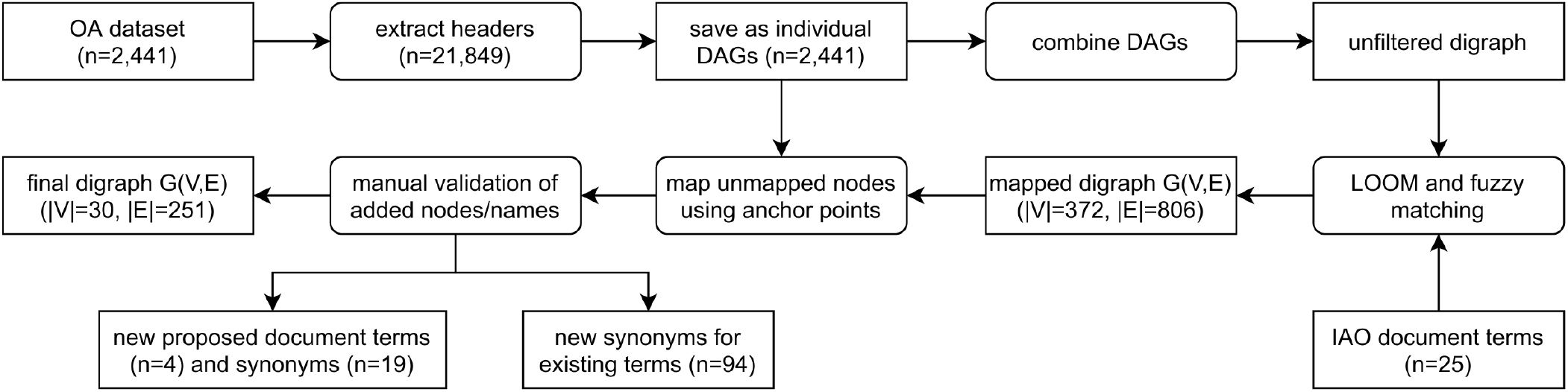
Flow diagram demonstrating the process of classifying publication sections with IAO terms.

**Fig. 4.**
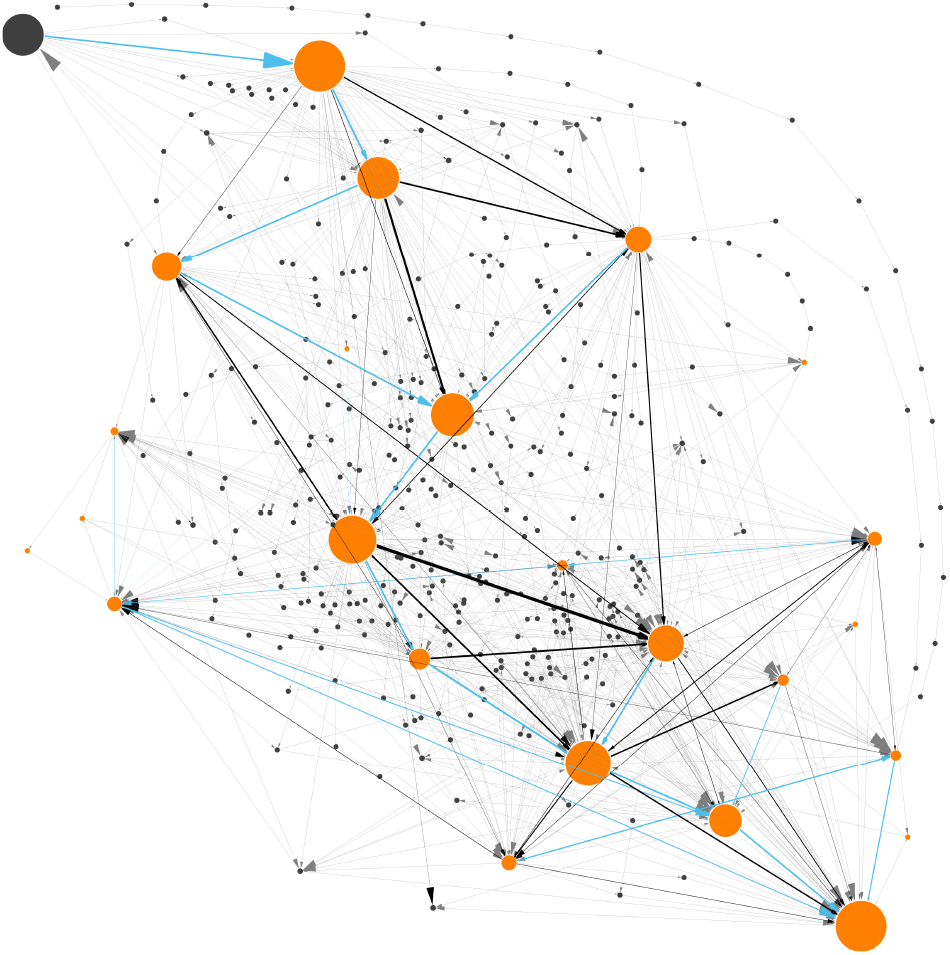
Digraph generated from analyzing section headers from 2,441 Open Access publications from PubMed Central. The digraph of the v2020-06-10 IAO model (17) consists of 372 unique nodes, of which 24 could be directly mapped to section terms (in orange) and the remainder are unmapped headers (in grey), and 806 directed edges. Relative node sizes and edge widths are directly proportional to the number of publications with these (subsequent) headers. Blue edges indicate the edge with the highest weight from the source node, edges that exist in fewer than 1% of publications are shown in light grey and the remainder in black.

However, after direct IAO mapping and fuzzy matching, unmapped headers still existed. To map these headings, we developed a new method using both the digraph and the individual DAGs. The headers are not repeated within a document/DAG, they are sequential and have a set order that can be exploited. Unmapped headers are assigned a section based on the digraph and the headers in the publication (DAG) that could be mapped (anchor headers), an example is given in Figure 5 where a header cannot be mapped to IAO terms. Auto-CORPus uses the LOOM, fuzzy matching and digraph prediction algorithms to annotate publication sections with IAO terms in the BioC full-text file.

**Fig. 5.**
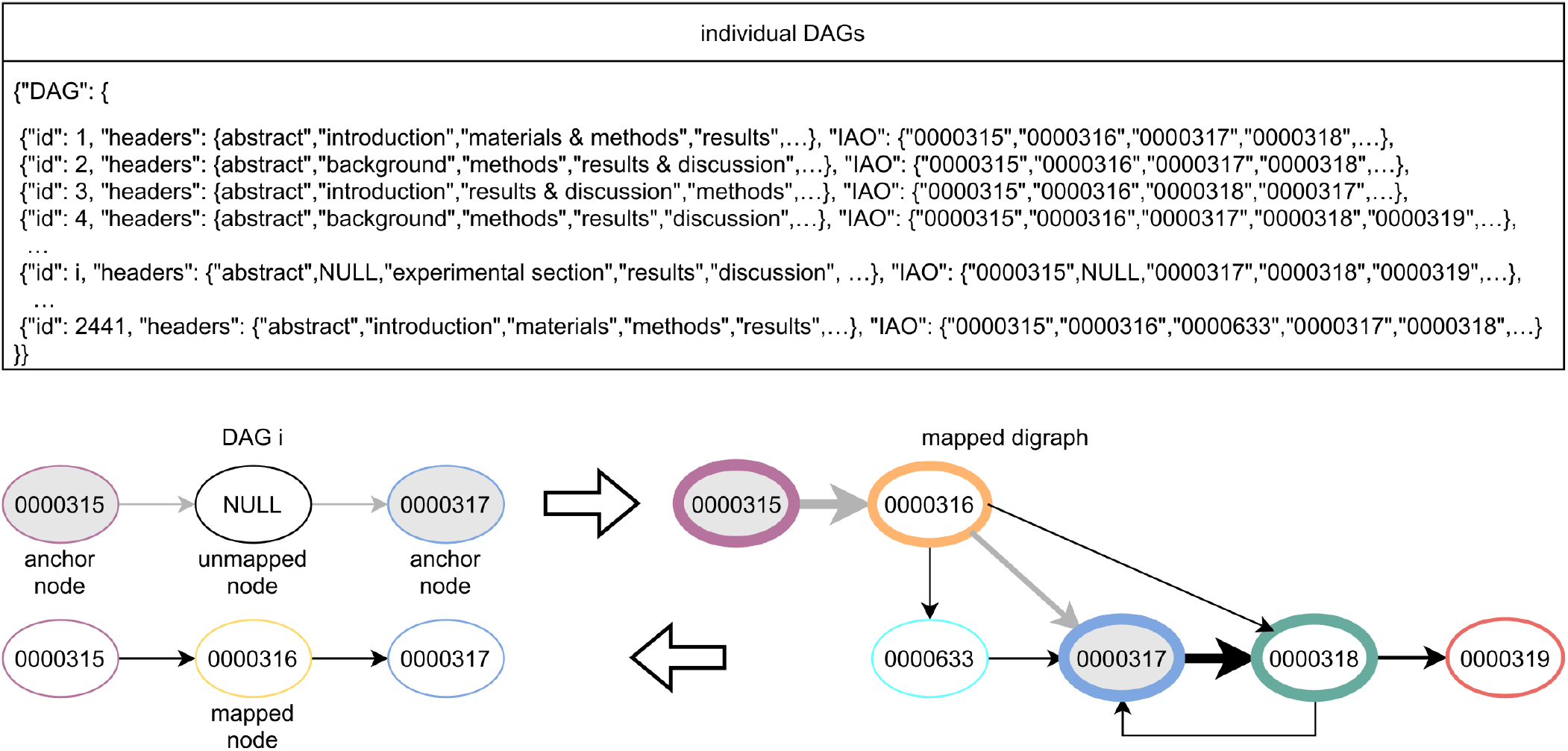
Pictographic example of using anchor nodes to map unmapped headers. Top panel illustrates the data structure of the individual directed acyclic graphs (DAGs) extracted from the OA dataset including the header and Information Artefact Ontology (IAO) term they are mapped to. The bottom panel indicates how anchor headers (nodes) are used to predict the IAO term of unmapped nodes based on the mapped digraph (see Figures 3 and 4) to find the shortest path between the mapped nodes and assigning unmapped nodes to nodes in the graph. Paths in the digraph between anchor nodes that are shorter than the DAG are mapped to the first anchor node.

#### New IAO terms and synonyms

We used the IAO classification algorithms to identify potential new IAO terms and synonyms. 348 headings from the OA dataset were mapped to IAO terms during the fuzzy matching or mapped based on the digraph using the publication structure and anchor headers. These headings were considered for inclusion in IAO as term synonyms. We manually evaluated each heading and Tables 2a and 2b list the 94 synonyms we identified for existing IAO terms.

**Table 2a.**
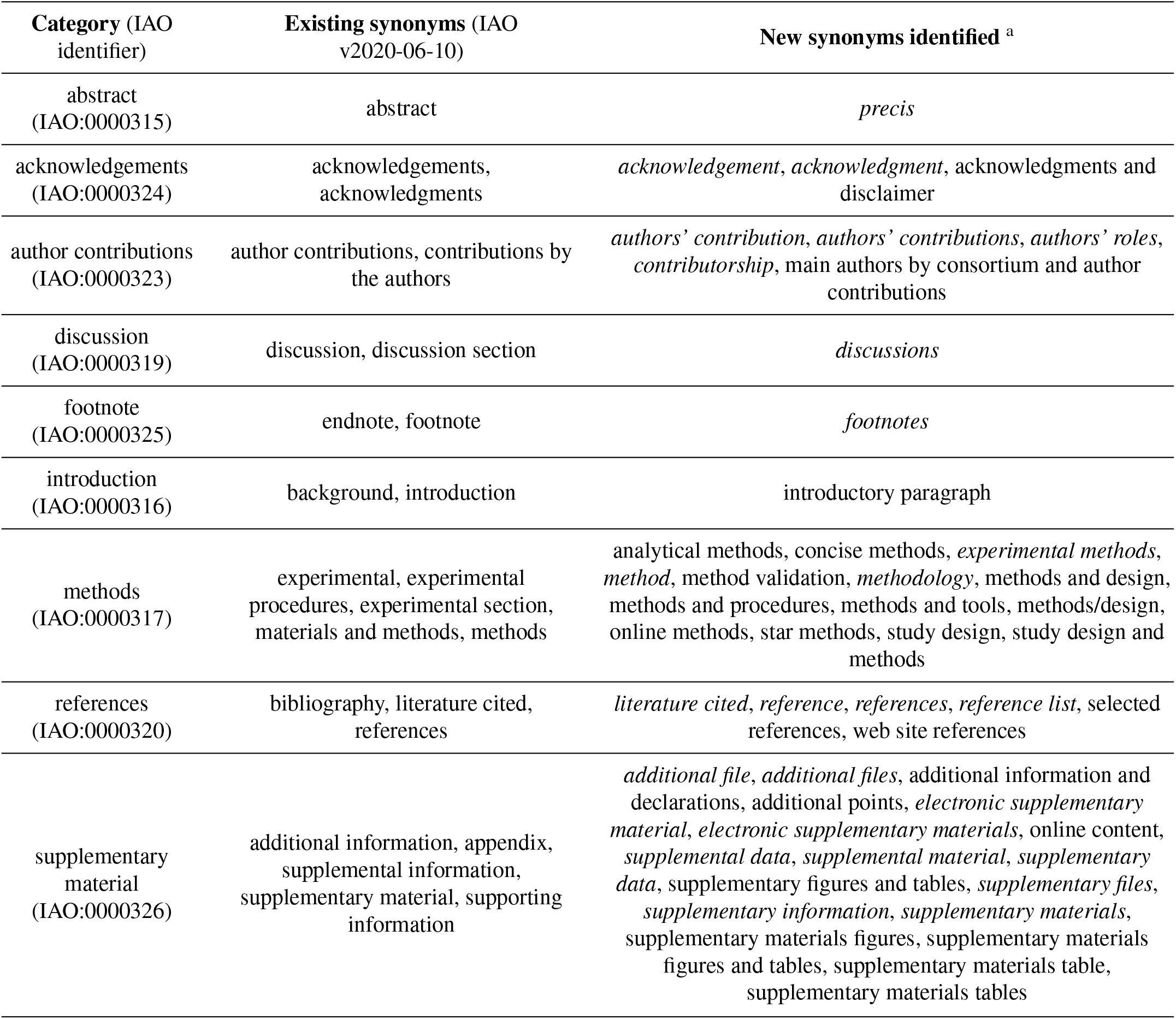
Newly identified synonyms for existing IAO terms (00003xx) from the digraph mapping of publications. Elements in *italics* have previously been submitted by us for inclusion into IAO and added in the v2020-12-09 release (18).

**Table 2b.** Newly identified synonyms for existing IAO terms (00006xx) from the digraph mapping of publications. Elements in *italics* have previously been submitted by us for inclusion into IAO and added in the v2020-12-09 release (18).

| Category (IAO identifier) | Existing synonyms (IAO v2020-06-10) | New synonyms identified <sup>a</sup> |
| --- | --- | --- |
| abbreviations (IAO:0000606) | abbreviations, abbreviations list, abbreviations used, list of abbreviations, list of abbreviations used | *abbreviation and acronyms, abbreviation list, abbreviations and acronyms, abbreviations used in this paper, definitions for abbreviations, glossary, key abbreviations, non-standard abbreviations, nonstandard abbreviations, nonstandard abbreviations and acronyms* |
| author information (IAO:0000607) | author information, authors' information | *biographies, contributor information* |
| availability (IAO:0000611) | availability, availability and requirements | *availability of data, availability of data and materials, data archiving, data availability, data availability statement, data sharing statement* |
| conclusion (IAO:0000615) | concluding remarks, conclusion, conclusions, findings, summary | conclusion and perspectives, summary and conclusion |
| conflict of interest (IAO:0000616) | competing interests, conflict of interest, conflict of interest statement, declaration of competing interests, disclosure of potential conflicts of interest | *authors' disclosures of potential conflicts of interest, competing financial interests, conflict of interests, conflicts of interest, declaration of competing interest, declaration of interest, declaration of interests, disclosure of conflict of interest, duality of interest, statement of interest* |
| consent (IAO:0000618) | consent | informed consent |
| ethical approval (IAO:0000620) | ethical approval | ethics approval and consent to participate, *ethical requirements, ethics, ethics statement* |
| funding source declaration (IAO:0000623) | funding, funding information, funding sources, funding statement, funding/support, source of funding, sources of funding | *financial support, grants, role of the funding source, study funding* |
| future directions (IAO:0000625) | future challenges, future considerations, future developments, future directions, future outlook, future perspectives, future plans, future prospects, future research, future research directions, future studies, future work | *outlook* |
| materials (IAO:0000633) | materials | data, data description |
| statistical analysis (IAO:0000644) | statistical analysis | statistical methods, statistical methods and analysis, statistics |
| study limitations (IAO:0000631) | limitations, study limitations | strengths and limitations, study strengths and limitations |

Digraph nodes that were not mapped to IAO terms but formed heavily weighted “ego-networks”, indicating the same heading was found in many publications, were manually evaluated for inclusion in IAO as new terms. For example, based on the digraph, we assigned *data* and *data description* to be synonyms of the *materials section*. The same process was applied to ego-networks from other nodes linked to existing IAO terms to add additional synonyms to simplify the digraph. Figure 6 shows the ego-network for *abstract*, and four main categories and one potential new synonym (*precis*, in red) were identified. From the further analysis of all ego-networks, four new potential terms were identified: *disclosure*, *graphical abstract*, *highlights* and *participants* – the latter is related to, but deemed distinct from, the existing *patients section* (IAO:0000635). Table 3 details the proposed definition and synonyms for these terms. The terms and synonyms described here will be submitted to the IAO, with our initial submission of one term and 59 synonyms accepted and included in IAO previously (v2020-12-09) (https://github.com/information-artifact-ontology/IAO/issues/234). Figure 7 shows the resulting digraph with only existing and newly proposed section terms. A major unmapped node is *associated data*, which is a header specific for PMC articles that appears at the beginning of each article before the abstract. In addition, IAO has separate definitions for *materials* (IAO:0000633), *methods* (IAO:0000317) and *statistical methods* (IAO:0000644) sections, hence they are separate nodes in the graph. The *introduction* is often followed by these headers to reflect the *methods section* (and synonyms), however there is also a major directed edge from *introduction* directly to *results* to account for *materials and methods* placed after the *discussion* and/or *conclusion* sections in some publications.

**Fig. 6.**
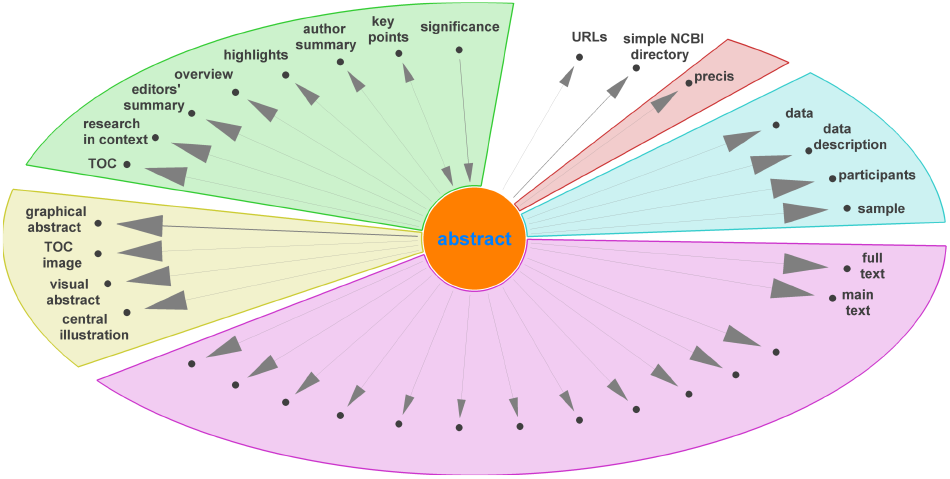
Unmapped nodes in the digraph (Figure 4) connected to ‘abstract’ as ego node, excluding corpus specific nodes, grouped into different categories. Unlabeled nodes are titles of paragraphs in the main text.

**Fig. 7.**
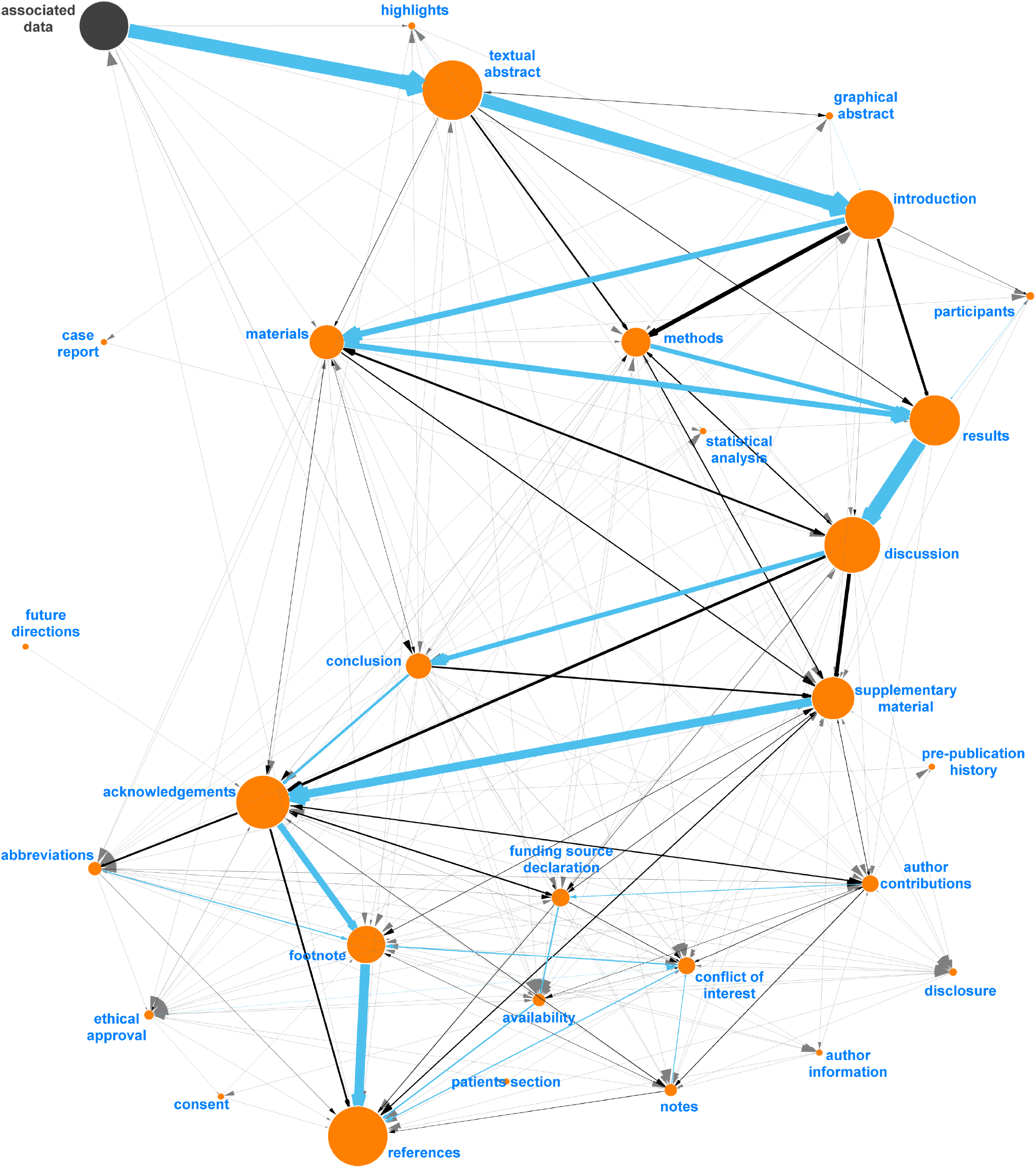
Final digraph model used in Auto-CORPus to classify paragraphs after fuzzy matching to IAO terms (v2020-06-10 (17)). This model includes new (proposed) section terms and each section contains new synonyms identified in this analysis. ‘Associated Data’ is included as this is a PMC-specific header found before abstracts and can be used to indicate the start of most articles, all IAO terms are indicated in orange.

**Table 3.** (A) Proposed new IAO terms to define publication sections that were derived from analysing the sections of 2,441 publications. (B) Proposed new IAO terms to define parts of a table section. ^a^ Elements in italics have previously been submitted by us for inclusion into IAO and added in the v2020-12-09 IAO release (18).

| (a) Proposed category | Proposed definition | Proposed synonyms <sup>a</sup> |
| --- | --- | --- |
| disclosure | “A part of a document used to disclose any associations by authors that might be perceived as to potentially interfere with or prevent them from reporting research with complete objectivity.” | author disclosure statement, declarations, disclosure, disclosure statement, disclosures |
| *graphical abstract* | *“An abstract that is a pictorial summary of the main findings described in a document.”* | central illustration, *graphical abstract*, TOC image, *visual abstract* |
| highlights | “A short collection of key messages that describe the core findings and essence of the article in concise form. It is distinct and separate from the abstract and only conveys the results and concept of a study. It is devoid of jargon, acronyms and abbreviations and targeted at a broader, non-technical audience.” | author summary, editors’ summary, highlights, key points, overview, research in context, significance, TOC |
| participants | “A section describing the recruitment of subjects into a research study. This section is distinct from the ‘patients’ section and mostly focusses on healthy volunteers.” | participants, sample |
| (b) Proposed category | Proposed definition | Proposed synonyms |
| table title | “A textual entity that names a table.” |  |
| table caption | “A textual entity that describes a table.” |  |
| table footer | “A part of a table that provides additional information about a specific other part of the table. Footers are spatially segregated from the rest of the table and are usually indicated by a superscripted number or letter, or a special typographic character such as †.” | table key, table note, table notes |

### Algorithms for processing tables

#### Auto-CORPus table JSON design

The BioC format does not specify how table content should be structured, leaving this open to the interpretation of implementers. For example, the PMC BioC JSON output describes table content using PMC XML (see the “pmc.key” file at https://ftp.ncbi.nlm.nih.gov/pub/wilbur/BioC-PMC/pmc.key). Including markup language within JSON objects presents data parsing challenges and interoperability barriers with non-PMC table data representations. We developed a simple table JSON format that is agnostic to the publication table source, can store multi-dimensional table content from complex table structures, and applies BioC design principles (5) to enable the annotation of entities and relations between entities. The table JSON stores table metadata of title, caption and footer. The table content is stored as “column headers” and “data rows”. The format supports the use of IAO to define the table metadata and content sections, however additional IAO terms are required to define table metadata document parts. Table 3 includes the proposed definition and synonyms for these terms. To compensate for currently absent IAO terms, we have defined three section type labels: *table title*, *table caption* and *table footer*. To support the text mining of tables, each column header and data row cell has an identifier that can be used to identify entities in annotations. Tables can be arranged into subsections, thus the table JSON represents this and includes subsection headings. Figure 8 gives an example of table metadata and content stored in the Auto-CORPus table JSON format. In addition to the Auto-CORPus key files, we make a table JSON schema available for the validation of table JSON files and to facilitate the use of the format in text analytics software and pipelines.

**Fig. 8.**
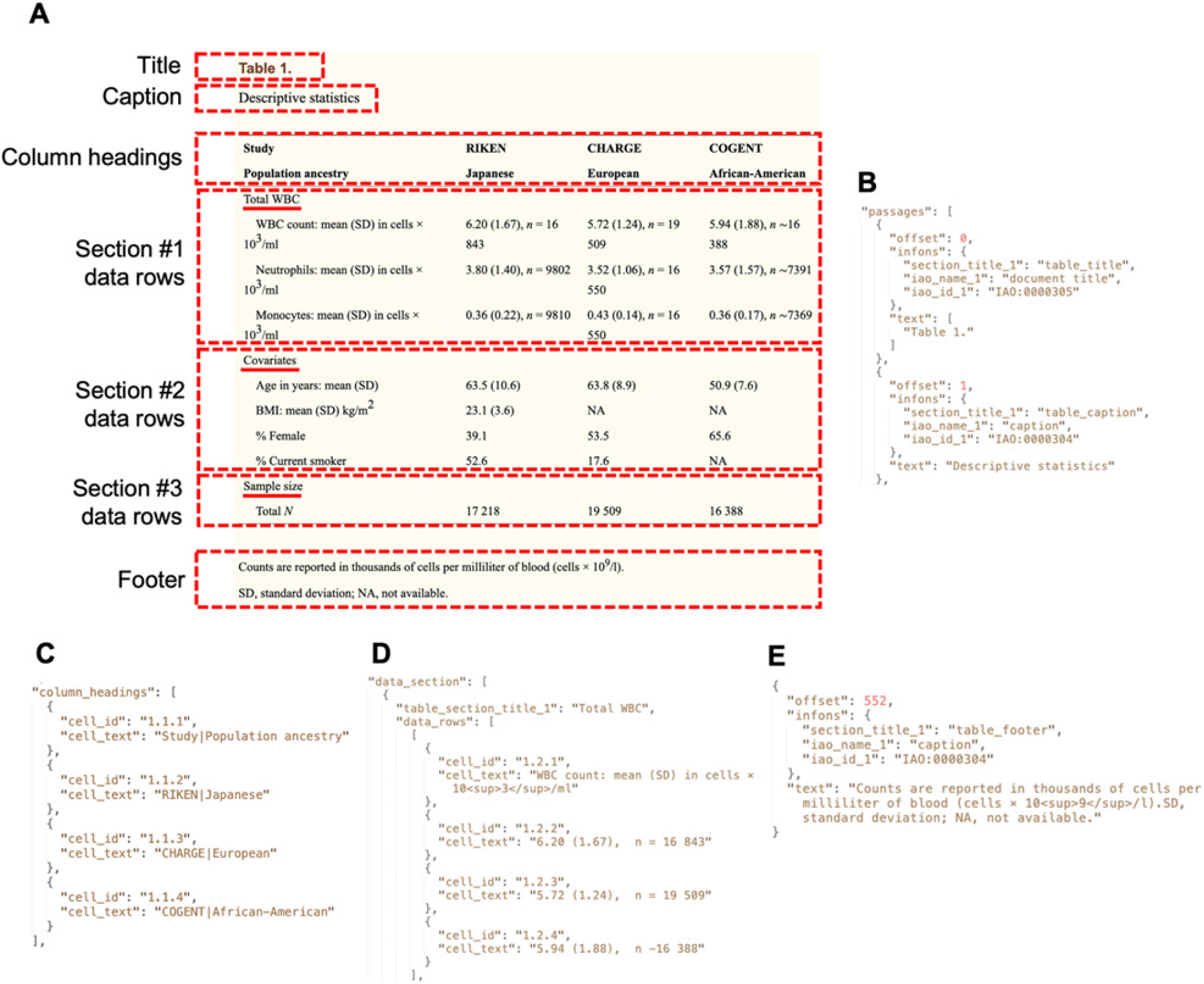
Extracts of the Auto-CORPus table JSON file generated to store metadata and content for an example table. (A) The parts of a table stored in table JSON. The section titles are underlined. The table shown is the PMC version (PMC4245044) of Table 1 from (19). (B) The title and caption table metadata stored in table JSON. (C) Each column heading in the table content is split between two rows, so the strings from both cells are concatenated with a pipe symbol in the table JSON. Headers that span multiple columns of sub-headers are replicated in each header cell as here with the pipe symbol. (D) The table content for the first row from the first section is shown in table JSON. Superscript characters are identified using HTML markup. (E) The footer table metadata stored in table JSON. The “autocorpus_tables.key” file describes the contents of the tables JSON file (https://github.com/omicsNLP/Auto-CORPus/blob/main/keyFiles/autocorpus_tables.key).

#### Processing table HTML

Tables can used within HTML documents for formatting web page layouts and are distinct from the *data tables* processed by Auto-CORPus. The configuration file set by the user identifies the HTML elements used to define data table containers, which include title, caption, footer and table content. The files processed can either be a full-text HTML file for inline tables and/or separate HTML files for individual linked tables. The Auto-CORPus algorithm for processing tables is based on the functional and structural table analysis method described by Milosevic and colleagues (8). The cells that contain navigational information such as column headers and section headings are identified. If a column has header strings contained in cells spanning multiple rows, the strings are concatenated with a pipe character separator to form a single column header string. The “super row” is a single text string that spans a complete row (multiple columns) within the table body. The “index column” is a single text string in the first column (sometimes known as a stub) within the table body when either only the first column does not have a header, or the cell spans more than one row. The presence of a super row or index column indicates a table section division where the previous section (if present) ends, and a new section starts. The super row or index column text string provides the section name. A nested array data structure of table content is built to relate column headers to data rows, working from top to bottom and left to right, with section headings occurring in between and grouping data rows. The algorithm extracts the table metadata of title, footer and caption. Table content and metadata are output in the table JSON format. The contents of table cells can be either string or number data types (we consider “true” and “false” booleans as strings) and are represented in the output file using the respective JSON data type. Cells that contain only scientific notation are converted to exponential notation and stored as a JSON number data type. All HTML text formatting is removed, however this distorts the meaning of positive exponents in text strings, for example *n=10^3^* is represented as *n=10^3^*. To preserve the meaning of exponents within text strings, superscript characters are identified using superscript HTML element markup, for example *n=10^3^*.

Some publication tables contain content that could be represented in two or more separate tables. These multi-dimensional tables use the same gridlines, but new column headers are declared after initial column headers and data rows have appeared in the table. New column headers are identified by looking down columns and classifying each cell as one of three types: numerical, textual, and a mix of numbers and texts. The type for a column is determined by the dominant cell type of all rows in a column excluding super rows. After the type of all columns are determined, the algorithm loops through all rows except super rows, and if more than half of cells in the row do not match with the columns’ types, the row is identified as a new header row, and the rows that follow the new headers are then regarded as a sub-table. Auto-CORPus represents sub-tables as distinct tables in the table JSON, with identical metadata to the initial table. Tables are identified by the table number used in the publication, so since sub-tables will share their table number with the initial table, a new identifier is created for sub-tables with the initial table number, an underscore, then a sub-table number such as “1_1”.

#### Processing table images - cell detection

Our current work includes the development of a pipeline of image processing operations to convert table images in JPG or PNG formats to table JSON. The three stages of the pipeline are cell detection, text recognition and table structure analysis.

The OpenCV library (https://github.com/opencv/opencv/tree/4.5.3) is used to preprocess images prior to cell detection. Image binarization is used to remove background colours, font colours and provide a high contrast between the target text and the background. Line detection is used to remove all grid lines from the table image, with the OpenCV minimum line length parameter configured to distinguish between table grid lines and short line characters such as “i”, capitalized “I” or “l”. Mathematical morphology processing is then used to identify cell text by conducting quantitative description and analysis of geometric shapes and structures based on set theory. Text blocks of interest are processed by thickening the black texts and connecting the discontinuous parts close to each other to blur black pixel regions, creating black “smudges” that identify the location of each text block. Rectangular boxes are drawn around each block to create individual cells that overlap with the original table cells and their spatial positions are indexed by the rectangle boundary’s horizontal and vertical coordinates. Figure 9 illustrates the cell detection steps.

**Fig. 9.**
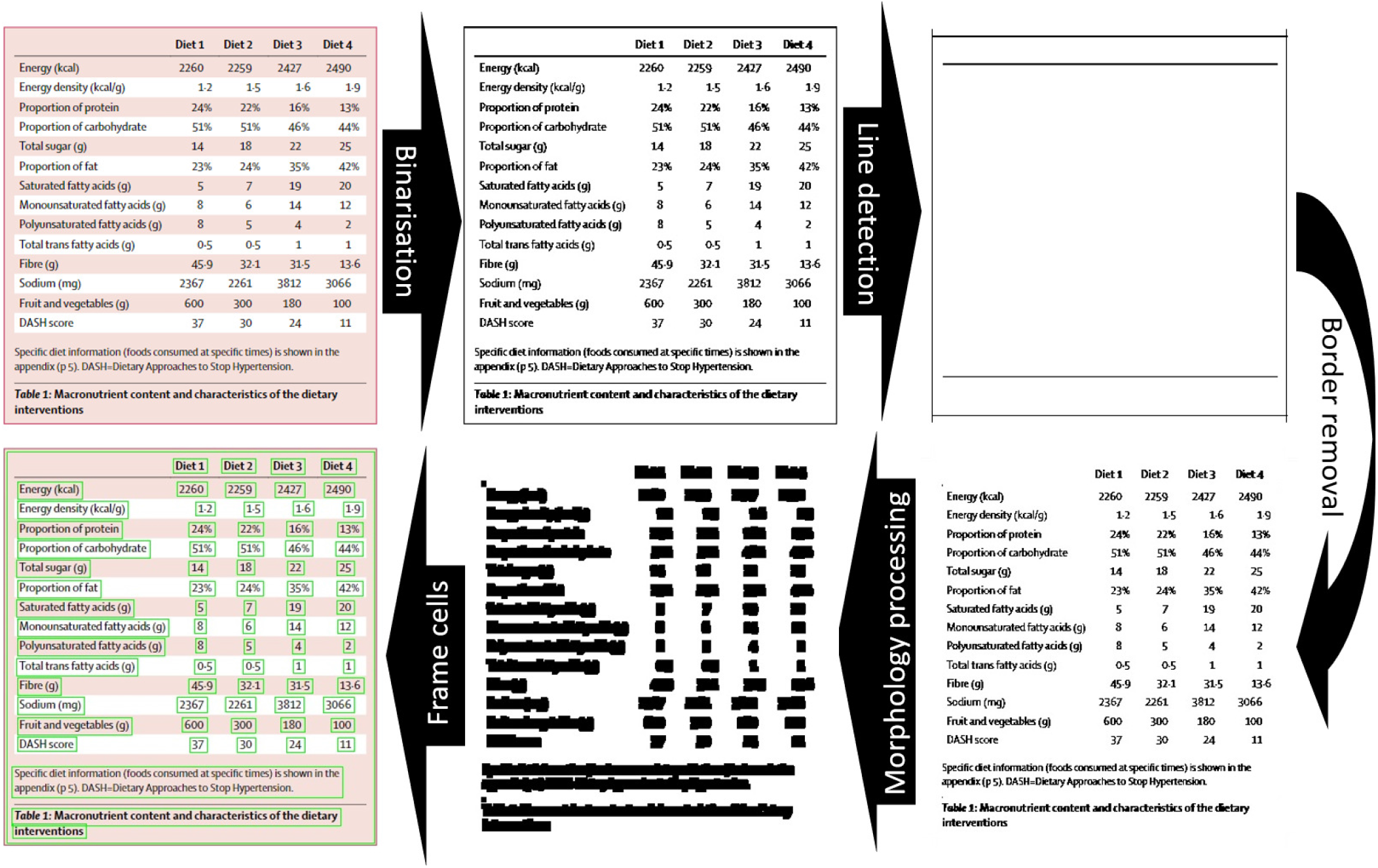
The image processing steps involved in cell detection. Binarization removes colour shading from table images to represent tables with black and white pixels only. Line detection is followed by the removal of horizontal and vertical grid lines in the image to reduce noise that could affect the morphology process. Text blocks are represented as blackened “smudges” to aid recognition, and finally, all cells containing text are identified and framed. The table is reproduced here under a Creative Common license from (20).

#### Processing table images - text recognition

Google’s Tesseract optical character recognition (OCR) engine is used through the Python-tesseract wrapper (https://pypi.org/project/pytesseract/) to recognise and extract text from the preprocessed table images. Tesseract is based on Long Short-Term Memory (LSTM) neural networks which enables users to train models aligned to their use cases. PyTesseract provides more than 100 trained libraries in different languages and we found that using the PyTesseract English language “eng.traineddata” library, the OCR engine correctly recognised 53.14% of cell text from the images in the publisher dataset (33,273 cells). Cells containing special characters such as superscripts, subscripts and Greek characters were rarely recognised. We trained a dataset using biomedical data to fine-tune the PyTesseract model, resulting in the OCR engine correctly recognising 87.92% (median, interquartile range: 86.14-90.58%) of cell text from the table images in the publisher dataset. Superscript and subscript characters were present in most cells not recognised.

#### Processing table images - table structure analysis

The text from each cell is analysed to determine the cell function and generate the structured table JSON output. The algorithm reads all of the cells line by line, from top-left to bottom-right, and creates a map of their relative positions. Spatial positioning and regular expression rules are then used to identify the table identifier, title, caption, footer and column headers. Super rows and index columns are identified and recognised as table section divisions (the end of a previous section and start of new section) and the cell text is included as the section title in the table JSON.

### Named-entity recognition on GWAS publications

To demonstrate the potential applicability of Auto-CORPus for text mining of the biomedical literature we used the GWAS publications from the OA dataset. This subset contains 1,200 publications that are present in the GWAS Central database (https://www.gwascentral.org/) which allows cross-referencing with manually extracted entities. Assigned randomly, 700 publications are used as a training set and 500 as a test set.

Prior work on GWAS publications from ourselves (unpublished), and from others (21), has shown phenotypes, P-values and single nucleotide polymorphisms (SNPs) can be extracted using regular expression matching. Here we focus on 5 different branches of important information that can be extracted about each study: platform recognition (company names), total number of SNPs assayed (total SNPs), sequencing technology used (exact array/assays), how quality control (QC) was performed (presence of QC, software version number), and whether imputation was performed (imputation including possible negation). The training set was created using data from GWAS Central which was manually curated (platform (list of strings), total SNPs passing QC (numbers in different formats) and imputation (binary)) and used for annotation, as well as a manually created list of entities for each of the categories. The sequencing technology (exact assay) was annotated by combining a search for the platform name, specific words (array, chip, etc.) with a regular expression algorithm to look for combinations of letters and numbers in sequence. For each of these branches, sentences containing relevant entities for training were extracted from the Auto-CORPus BioC JSON output to create a training set for each branch, for this we restricted ourselves to the (statistical) methods/materials sections only (IAO:0000317, IAO:0000633, and IAO:0000644).

SpaCy 3.0 (https://spacy.io/models/enen_core_web_lg) was used to develop each of the algorithms using word embedding and a multilayer convolution neural network (CNN) along with residual connections (22) and models were optimized for accuracy (over speed). The maximum batch size was set to 1000 and the proprietary spaCy quickstart configuration file was deployed for every single branch model, allowing for unlimited epochs until the model scoring plateaus.

### Comparative analysis of outputs

The correspondence between PMC BioC and Auto-CORPus BioC outputs were compared to evaluate whether all information present in the PMC BioC output also appears in the Auto-CORPus BioC output. This was done by analysing the number of characters in the PMC BioC JSON that appear in the same order in the Auto-CORPus BioC JSON using the longest common sub-sequence method. With this method, overlapping sequences of characters that vary in length are extracted from the PMC BioC string to find a matching sequence in the Auto-CORPus string. With this method it can occur that a subsequence from the PMC BioC matches to multiple parts of the Auto-CORPus BioC string (e.g. repeated words). This is mitigated by evaluating matches of overlapping/adjacent subsequences which should all be close to each other as they appear in the PMC BioC text.

This longest common subsequence method was applied to each individual paragraph of the PMC BioC input and compared with the Auto-CORPus BioC paragraphs. This method was chosen over other string metric algorithms, such as the Levenshtein distance or cosine-similarity, due to it being non-symmetric/unidirectional (the Auto-CORPus BioC output strings contain more information (e.g., figure/table links, references) than the PMC BioC output) and ability to directly extract different characters.

## Results

### Data for the evaluation of algorithms

We attempted to download PMC BioC JSON format for all 1,200 GWAS PMC publications in our OA dataset, but only 766 were available as BioC from the NCBI server. We refer to this as the “PMC BioC dataset”. For the 766 PMC articles we could obtain a NCBI BioC file for, we processed the equivalent PMC HTML files using Auto-CORPus. We used only the BioC output files and refer to this as the “Auto-CORPus BioC dataset”. To compare the Auto-CORPus BioC and table outputs for PMC and publisher-specific versions, we accessed 163 Nature Communication and 5 Nature Genetics articles that overlap with the OA dataset and were not present in the publisher dataset, so they were unseen data. These journals have linked tables, so full-text and all linked table HTML files were accessed (367 linked table files). Auto-CORPus configuration files were setup for the journals to process the publisher-specific files and the BioC and table JSON output files were collated into what we refer to as the “linked table dataset”. The equivalent PMC HTML files from the OA dataset were also processed by Auto-CORPus and the BioC and table JSON files form the “inline table dataset”.

### Performance of Auto-CORPus full-text processing

The proportion of characters from 3,195 full-text paragraphs in the PMC BioC dataset that also appear in the Auto-CORPus BioC dataset in the same order in the paragraph string were evaluated using the longest common subsequence method. The median and interquartile range of the (left-skewed) similarity are 100% and 100-100%, respectively. Differences between the Auto-CORPus and PMC outputs are shown in Table 4 and relate to how display items, abbreviations and links are stored, and different character encodings. A structural difference between the two outputs is in how section titles are associated to passage text. In PMC BioC the section titles (and subtitles) are distinct from the passages they describe as both are treated as equivalent text. The section title occurs once in the file and the passage(s) it refers to follows it. In Auto-CORPus BioC the (first level) section titles (and subtitles) are linked directly with the passage text they refer to, and are included for each paragraph. Auto-CORPus uses IAO to classify text sections so, for example, the introduction title and text are grouped into a section annotated as introduction, rather than splitting these into two subsections (introduction title and introduction text as separate entities in the PMC BioC output) which would not fit with the IAO structure.

**Table 4.**
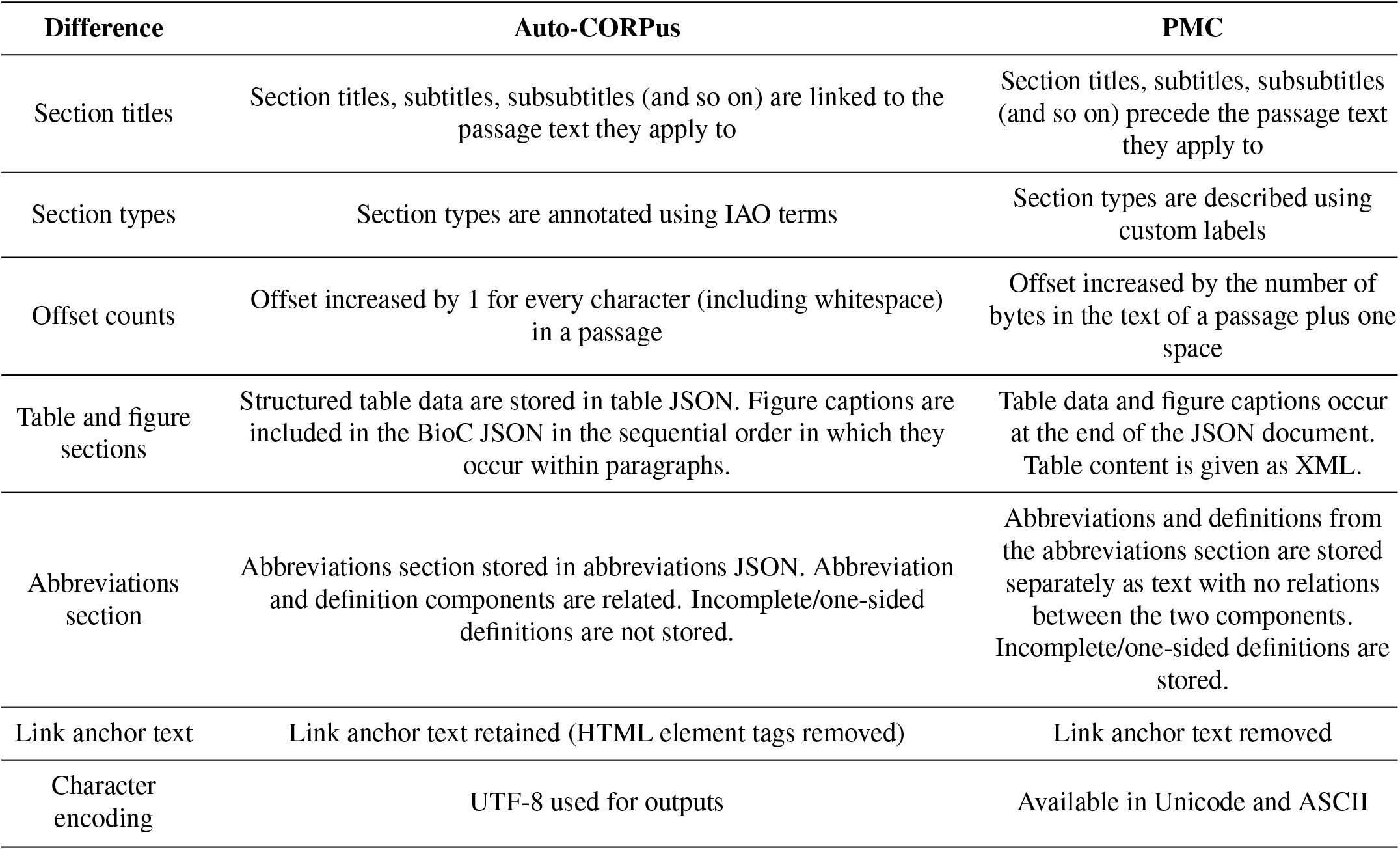
Differences between the Auto-CORPus BioC and PMC BioC JSON outputs.

The Auto-CORPus BioC output includes the figure captions where they appear in the text and a separate table JSON file to store the table data, whereas the PMC BioC adds these data at the end of the JSON document and provides table content as a block of XML. Abbreviation sections are not included in the Auto-CORPus BioC output since Auto-CORPus provides a dedicated abbreviations JSON output. In the PMC BioC format the abbreviations and definitions are not related, whereas in the Auto-CORPus abbreviations JSON output the two elements are related. If an abbreviation does not contain a definition in the abbreviations section (perhaps due to an editorial error), PMC BioC will include the undefined thus meaningless abbreviation string, whereas Auto-CORPus will ignore it. Link anchor text to figures, tables, references and URLs are retained in the Auto-CORPus output but removed in the PMC BioC output. The most common differences between the two BioC versions is the encodings/strings used to reflect different whitespace characters and other special characters, with the remaining content being identical.

Last, the proportion of characters from 9,468 full-text paragraphs in the publisher dataset that also appear in the Auto-CORPus PMC BioC dataset in the same order in the paragraph string were evaluated. The median and interquartile range of the (left-skewed) similarity is also 100% and 100-100%, respectively, and differences between the PMC and publisher-versions also relate to the same as reported in Table 4.

#### Performance of Auto-CORPus table processing

We assessed the accuracy of the table JSON output generated from non-PMC linked tables compared with table JSON output generated from the equivalent PMC HTML with inline tables. The comparative analysis method described above was used for comparing BioC output from the linked table and inline table datasets, except here it was applied to both strings (bidirectional, taking the maximum value of both outcomes). This is equivalent to the Levenshtein similarity applied to transform the larger string into the smaller string, with the exception that the different characters for both comparisons are retained for identifying the differences. The correspondence between table JSON files in the linked table and inline table datasets was calculated as the number of characters correctly represented in the publishers table JSON output relative to the PMC versions (also using the (symmetric) longest common subsequence method). Both the text and table similarity are represented as the median (inter-quartile range) to account for non-normal distributions of the data. Any differences identified during these analyses were at the paragraph or table row level, enabling manual investigation of these sections in a side-by-side comparison of the files.

The proportion of characters from 367 tables in the linked table dataset that also appear in the inline table dataset in the same order in the cell or text string were evaluated. The median and interquartile range of the (left-skewed) similarity is 100% and 99.79-100.00%, respectively. We found that there were structural differences between some of the output files where additional data rows were present in the JSON files generated from the publisher’s files. This occurred because cell value strings in tables from the publisher’s files were split across two rows, however in the PMC version the string was formatted (wrapped) to be contained within a single row. The use of different table structures to contain the same data resulted in accurate but differing table JSON outputs. Most of the differences between table content and metadata values pertain to the character encoding used in the different table versions. For example, we have found different uses of hyphen/em dash/en dash/minus symbols between different versions, and Greek letters were represented differently in the different table versions. Other differences are related to how numbers are represented in scientific notation. If a cell contains a number only, then it is represented as a JSON number data type in the output. However, if the cell contains nonnumeric characters, then there is no standardisation of the cell text and the notation used (e.g., the × symbol or E notation) will be reproduced in the JSON output. When there is variation in notation between sources, the JSON outputs will differ. Other editorial differences include whether thousands are represented with or without commas and how whitespace characters are used. Despite these variations there was no information loss between processed inline and linked tables.

### Application of NER on GWAS publications

We used the BioC JSON output from Auto-CORPus to filter out sentences in the methods sections that contain information on the platforms, assays, total number of genetic variants, quality control and imputation that were used (Supplementary Methods). We trained five separate algorithms for named-entity recognition (NER) using 700 GWAS publications and evaluated these on 500 GWAS publications of the test set. The F1scores for the five tasks are between 0.82-1.00 (Table 5) with examples of the pipeline output (after merging output for each branch) for different sentences from the test set given in Figure 10. The branches with the best performance were the platform, quality control and imputation entities (all F1scores over 0.95). The least successful was the algorithm to recognise the exact assays used with an F1-score of 0.82.

**Fig. 10.**
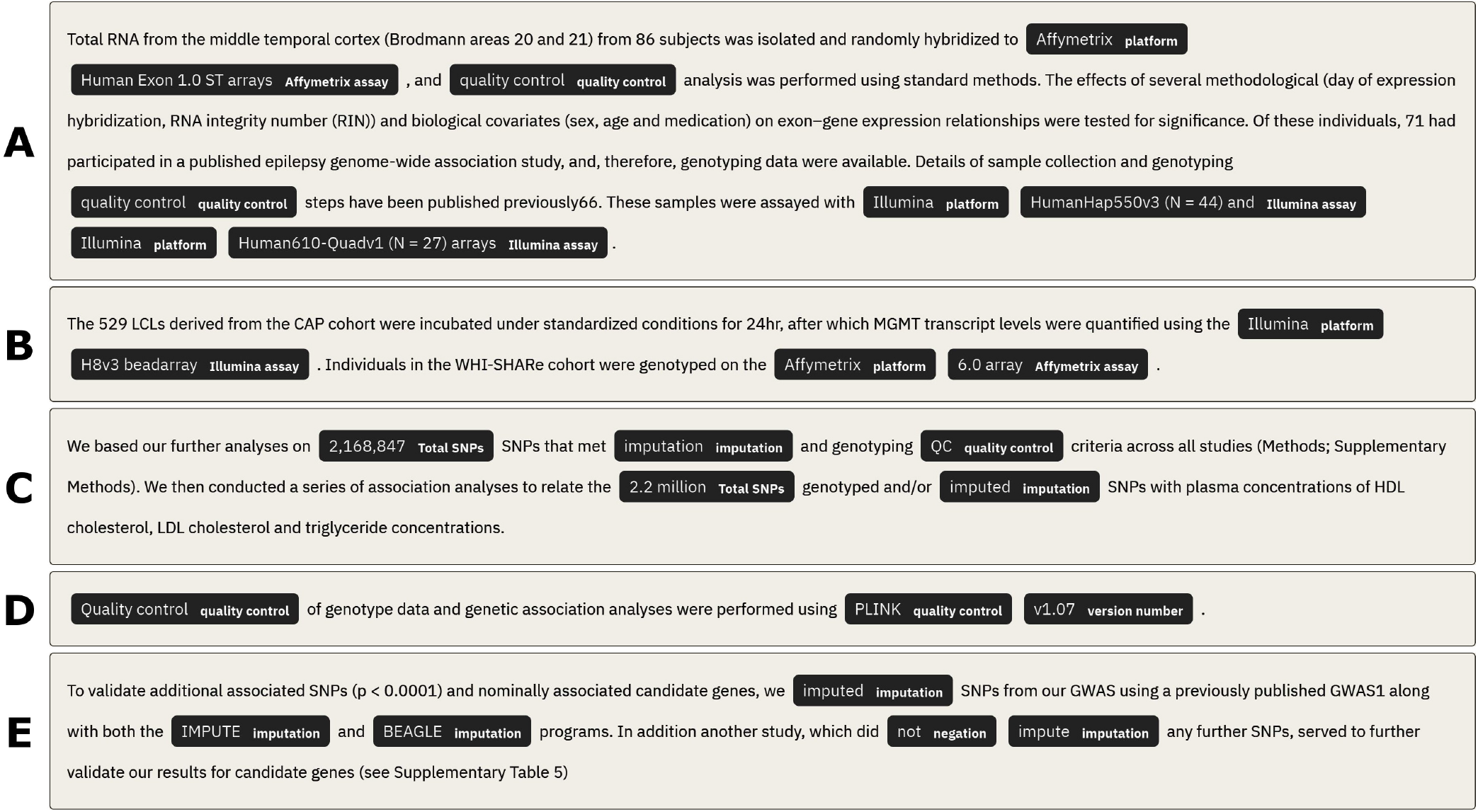
Example sentences from the GWAS publications test set (n=500) with recognized entities tagged. These examples combine the output from each branch for the individual NER tasks. (A) Details recognised platforms (platform), sequencing technology used (Affymetrix assay, Illumina assay), and presence of quality control (quality control). The latter 4 entities recognized were found as a ‘single’ Illumina assay but split after post-processing and combining with the platform NER algorithm output. (B) Details recognised platforms (platform) and sequencing technology used (Affymetrix assay, Illumina assay). (C) Details total number of SNPs (Total SNPs), whether imputation was used (imputation), and presence of quality control (quality control). (D) Details presence of quality control (quality control, version number). (E) Details whether imputation was used (imputation, negation). Note: software programs for quality control and imputation were labelled as the main category (quality control, imputation).

**Table 5.** F1-score for each of the 5 branches of GWAS NER evaluated on 500 publications from the test set. ^a^ The ‘no imputation’ tag is distinct from negation and focuses on words such as ‘unimputed’ and ‘non-imputed’, whereas negation focuses on entities such as ‘did not perform’ and ‘was not used’ that suggests ambiguity (i.e. relationship to concepts needs to be inferred). These tags are given as output of our pipeline (also see Figure 10).

| Branch (entity tags) | F1-score |
| --- | --- |
| Platform technology (platform) | 1.00 |
| Total number of SNPs (Total SNPs) | 0.89 |
| Assay name (Illumina assay, Affymetrix assay, Perlegen assay) | 0.82 |
| QC (quality control, negation, version number) | 0.98 |
| Imputation (imputation, negation, no imputation)<sup>a</sup> | 0.95 |

## Discussion

### Strengths and Limitations

We have shown that AutoCORPus brings together and bolsters several disjointed standards (BioC and IAO) and algorithmic components (for processing tables and abbreviations) of scientific literature analytics into a convenient and reliable tool for standardising full-text and tables. The BioC format is a useful but not ubiquitous standard for representing text and annotations. Auto-CORPus enables the transformation of the widely available HTML format into BioC JSON following the setup of a configuration file associated with the structure of the HTML documents. The use of the configuration file drives the flexibility of the package, but also restricts use to users who are confident exploring HTML document structures. We make available the configuration files used in th e evaluations described in this paper. To process additional sources, an upfront time investment is required from the user to explore the HTML structure and set the configuration file. We will be increasing the number of configuration files available for larger publishers, and we help non-technical users by providing documentation to explain how to setup configuration files. We welcome configuration files submitted by users and the documentation describes the process for users to submit files. Configuration files contain a section for tracking contributions made to the file, so the names of authors and editors can be logged. Once a configuration file has been submitted and tested, the file will be included within the Auto-CORPus package and the user credited (should they wish) with authorship of the file.

The inclusion of IAO terms within the Auto-CORPus BioC output standardises the description of publication sections across all processed sources. The digraph that is used to assign unmapped paragraph headers to standard IAO terms was constructed using both GWAS and MWAS literature to avoid training it to be used for a single domain only. We have tested the algorithms on PMC articles from three different physics (Frontiers in Physics, Nature Physics, Physical Review Letters) and three psychology (Frontiers in Psychology, Psychological Bulletin, Psychological Science) journals to confirm the BioC JSON output and IAO term recognition extend beyond only biomedical literature. The header terms from these articles were mapped to relevant IAO terms. Since ontologies are stable but not static, any resource or service that relies on one ontology structure could become outdated or redundant as the ontology is updated. We will rerun the fuzzy matching of headers to IAO terms and regenerate the digraph as new terms are introduced to the document part branch of IAO. We have experience of this when our first group of term suggestions based on the digraph were included into the IAO.

The BioC output of abbreviations contains the abbreviation, definition and the algorithm(s) by which each pair was identified. One limitation of the current full-text abbreviation algorithm is that it searches for abbreviations in brackets and therefore will not find abbreviations for which the definition is in brackets, or abbreviations that are defined without use of brackets. The current structure of the abbreviation JSON allows additional methods to be included to the two methods we currently use. Adding further algorithms to find different types of abbreviation in the full-text is considered as part of future work.

Auto-CORPus implements a method for extracting table structures and data that was developed to extract table information from XML formatted tables (8). The use of the configuration file for identifying table containers enables the table processing to be focused on relevant data tables and exclude other tables associated with web page formatting. Auto-CORPus is distinct from other work in this field that uses machine learning methods to classify the types of information within tables (23). Auto-CORPus table processing is agnostic to the extracted variables, with the only distinction made between numbers and strings for the pragmatic reason of correctly formatting the JSON data type. The table JSON files could be used in downstream analysis (and annotation) of cell information types, but the intention of Auto-CORPus is to provide the capability to generate a faithful standardised output from any HTML source file. We have shown high accuracy (>99%) for the tables we have processed with a configuration file and the machine learning method was shown to recover data from 86% of tables (23). Accurate extraction is possible across more data sources with the Auto-CORPus rule-based approach, but a greater investment in setup time is required.

Auto-CORPus focusses on HTML versions of articles as these are readily and widely available within the biomedical domain. Currently the processing of PDF documents is not supported, but the work by the Semantic Scholar group to convert PDF documents to HTML is encouraging as they observed that 87% of PDF documents processed showed little to no readability issues (4). The ability to leverage reliable document transformation will have implications for processing supplementary information files and broader scientific literature sources which are sometimes only available in PDF format, and therefore will require conversion to the accessible and reusable HTML format.

### Future Research and Conclusions

We found that the tables for some publications are made available as images, so could not be processed by Auto-CORPus. To overcome this gap in publication table standardisation, we are refining a plugin for Auto-CORPus that provides an algorithm for processing images of tables. The table image processing pipeline leverages Google’s Tesseract optical character recognition engine to extract text from preprocessed table images. During our preliminary evaluation of the plugin, it achieved an accuracy of 88% when processing a collection of 200 JPG and PNG table images taken from 23 different journals. Although encouraging, there are caveats in that the image formats must be of high resolution, the algorithm performs better on tables with gridlines than tables without gridlines, special characters are rarely interpreted correctly, and cell text formatting is lost. We are fine-tuning the Tesseract model by training new datasets on biomedical data. An alpha release of the table image processing plugin is available with the Auto-CORPus package.

The authors are involved in omics health data NLP projects that use Auto-CORPus within text mining pipelines to standardise and optimise biomedical literature ahead of entity and relation annotations and have given examples in the Supplementary Materials of how the Auto-CORPus output was used to train these algorithms. The BioC format supports the stand-off annotation of linguistic features such as tokens, part-of-speech tags and noun phrases, as well as the annotation of relations between these elements (5). We are developing machine learning methods to automatically extract genome-wide association study (GWAS) data from peer-reviewed literature and have given an example here of some of the tools that are being developed. High quality annotated datasets are required to develop and train NLP algorithms and validate the outputs. We are developing a GWAS corpus that can be used for this purpose using a semi-automated annotation method. The GWAS Central database is a comprehensive collection of summary-level GWAS findings imported from published research papers or submitted by study authors (13). For GWAS Central studies, we used Auto-CORPus to standardise the full-text publication text and tables. In an automatic annotation step, for each publication, all GWAS Central association data was retrieved. Association data consists of three related entities: a phenotype/disease description, genetic marker and an association P-value. A named entity recognition algorithm identifies the database entities in the Auto-CORPus BioC and table JSON files. The database entities and relations are mapped back onto the text, by expressing the annotations in BioC format and appending these to the relevant BioC element in the JSON files. The automatic annotations are then manually evaluated using the TeamTat text annotation tool which provides a user-friendly interface for annotating entities and relations (24). We use TeamTat to manually inspect the automatic annotations and modify or remove incorrect annotations, in addition to including new annotations that were not automatically generated. TeamTat accepts BioC input files and outputs in BioC format, thus the Auto-CORPus files that have been automatically annotated are suitable for importing into TeamTat. Work to create the GWAS corpus is ongoing, but the convenient semi-automatic process for creating high-quality annotations from biomedical literature HTML files described here could be adapted for creating other gold-standard corpora.

In related work, we are developing a corpus for MWAS for metabolite named-entity recognition to enable the development of new NLP tools to speed up literature review. As part of this, the active development focuses on extending Auto-CORPus to analyse preprint literature and supplementary materials, improving the abbreviation detection, and development of more configuration files. Our preliminary work on preprint literature has shown we can map paragraphs in Rxiv versions to paragraphs in the peer-reviewed manuscript with the high accuracy (average similarity of paragraphs >95%). The flexibility of the Auto-CORPus configuration file enables researchers to use Auto-CORPus to process publications and data from a broad variety of sources to create reusable corpora for many use cases in biomedical literature and other scientific fields.

## ACKNOWLEDGEMENTS

We thank Mohamed Ibrahim (University of Leicester) for identifying different configurations of tables for different HTML formats.

## AUTHOR CONTRIBUTIONS

TB and JMP designed and supervised the research and wrote the manuscript. TS developed the BioC outputs and led the coding integration aspects. YH developed the section header standardization algorithm and implemented the abbreviation recognition algorithm. ZL developed the table image recognition and processing algorithm. SS developed the table extraction algorithm and main configuration file. CP developed configuration files for preprint texts. NARM developed the NER algorithms for GWAS entity recognition. NARM, FM, CSY, ZL and CP tested the package and performed comparative analysis of outputs. TR refined standardization of full-texts and contributed algorithms for character set conversions. All authors read, edited and approved the manuscript.

